# Potentially repurposable drugs for schizophrenia identified from its interactome

**DOI:** 10.1101/442640

**Authors:** Kalyani B. Karunakaran, Srilakshmi Chaparala, Madhavi K. Ganapathiraju

**Affiliations:** Supercomputer Education and Research Centre, Indian Institute of Science; Department of Biomedical Informatics, University of Pittsburgh; intelligent Systems Program, University of Pittsburgh

## Abstract

From the schizophrenia drug-target interactome,^1^ we studied the drugs that targeted multiple proteins in the interactome, or those that target proteins with many targets, or those that target novel (computationally predicted) interactors of schizophrenia associated proteins. In schizophrenia, gene expression has been described as a measurable aspect of the disease reflecting the action of risk genes. We studied each of the selected drugs using the NextBio software suite, and shortlisted those that had a negative correlation with gene expression of schizophrenia. This analysis resulted in 12 drugs whose differential gene expression (drug versus normal) had an anti-correlation with differential expression for schizophrenia (disorder versus normal). Some of these drugs were already being tested for their clinical activity in schizophrenia and other neuropsychiatric disorders. Several proteins in the protein interactome of the targets of several of these drugs were associated with various neuropsychiatric disorders. The network of genes which were differentially expressed on drug treatment, and had an anti-correlation with gene expression in schizophrenia, were significantly enriched in pathways relevant to schizophrenia etiology and GWAS genes associated with traits or diseases that had pathophysiological overlap with schizophrenia. Drugs that are structurally similar to the shortlisted drugs, or targeted the same genes as these drugs, have also demonstrated clinical activity in schizophrenia and other related disorders. This integrated computational analysis may help translate insights from the schizophrenia drug-protein interactome to clinical research - an important step, especially in the field of psychiatric drug development, facing a high failure rate.

## Introduction

Schizophrenia is a complex disorder with a cumulative impact of variable genetic effects coupled with environmental factors.^2^ The Schizophrenia Working Group of the Psychiatric Genomics Consortium (PGC) had identified 108 genetic loci that likely confer risk for schizophrenia.^3^ Prior to this, around 25 genes were being studied for their association with the disorder.^4^ While the role of genetics has been clearly validated by the genome-wide association studies (GWAS), the functional impact of the risk variants is not well understood. Several of the schizophrenia genes, especially those implicated by the GWAS have unknown functions and/or pathways. To discover the functional role of these genes, and promote discovery of novel therapeutics, we had carried out a computational analysis of protein-protein interactions (PPI) network, or the interactome, of schizophrenia associated genes.^1^ The schizophrenia interactome, comprising of 101 schizophrenia genes and about 1,900 PPIs, provided valuable results highlighting the functions and pathways tied to schizophrenia genes through their PPIs.^1^ A valuable result from this study was the drug-target interactome that showed various drugs that target proteins in the schizophrenia interactome. Many of these drugs were labeled for therapeutic value for nervous system as was expected, but there were also several other drugs that were labeled for other anatomical systems in the human body.

As drug approvals for psychiatric indications have been facing a high failure rate in the last few years,^5^ it would be beneficial to study whether these drugs that target proteins from the schizophrenia interactome could be repurposed for treatment of schizophrenia. Finding alternate uses for drugs that have been approved and are in use for other indications would be optimal, and such uses are being found in recent years.^6–8^

Diseases are often driven by an abnormal or perturbed expression of a multitude of genes which together constitute unique differential (gene) expression signatures (DES).^9^ Drugs administered to treat these diseases often revert the perturbed expression of these genes to their normal levels of expression. DES under disease and non-disease conditions are quantified using RNA sequencing methods based on microarrays and next-generation sequencing and are deposited in online repositories, which make the data freely available for integrated computational analyses.^10^ Similarly, DES corresponding to a number of drugs is made available through Connectivity Map.^11^ In order to analyze the suitability of these drugs for repurposing, we build over the results from our previous work on schizophrenia interactome discovery and analysis,^1^ utilizing the large transcriptomic databases, and employing a bioinformatics software suite named BaseSpace Correlation Engine.^12^

Many genetic variants associated with schizophrenia, viz. susceptibility genes such as DTNBP1, DAOA, NRG1 and RGS4, show differential gene expression in post-mortem brain samples obtained from schizophrenia patients compared with normal controls.^13^ In schizophrenia, it has been pointed out that the effect of genetic variants may, in fact, be reflected on gene expression rather than on the structure of the proteins coded by these genes. As a result of this, DES has been described as a psychiatric endophenotype which is a measurable aspect of the disease reflecting the action of the susceptibility genes.^14^ So, our method to identify repurposable drugs may be tested on schizophrenia, in which differential gene expression plays a critical role. We pruned the large list of drugs obtained from the schizophrenia drug-protein interactome using our in silico protocol, and present here a shortlist of drugs that may potentially be repurposed for schizophrenia.

## Results

In our prior work,^1^ we presented a few hundred drugs that target any of the proteins in the Schizophrenia Interactome^1^. Here, we analyzed these drugs to identify those that can potentially be repurposable for schizophrenia. We carried out bioinformatics analysis on these drugs to answer the following questions: Do these drugs induce a differential expression that compensates the different expression due to the disease? Have any of these drugs been considered for clinical trials? Are the genes that are targeted by these drugs or whose expression is altered by these drugs involved in biological pathways or processes relevant to schizophrenia?

### Bioinformatics analysis

We followed an established approach to identify drugs that have opposite differential expression to the differential expression of schizophrenia (i.e., genes over-expressed in schizophrenia are under-expressed by drug treatment and vice versa).^9^ We identified such drugs using the BaseSpace Correlation Engine software suite.^12^ This analysis resulted in 12 drugs whose differential gene expression (drug versus normal) had an anti-correlation with differential expression for schizophrenia (disorder versus normal). Although in each case, there are some genes that are differentially expressed in the same direction for both the drug and disorder, the overall effect on the entire transcriptome has an anti-correlation, leading to 12 drugs as potential candidates for further studies (Table 1, Figure 1). The top 5 drugs by the score of anticorrelation are cromoglicic acid, bepridil, acetazolamide, dimenhydrinate, cinnarizine, of which bepridil and dimenhydrinate may be excluded due to their side-effects related to nervousness and hallucinations (see Table 1), thus leaving cromoglicic acid, acetazolamide and cinnarizine as top candidates.

**Table 1.**
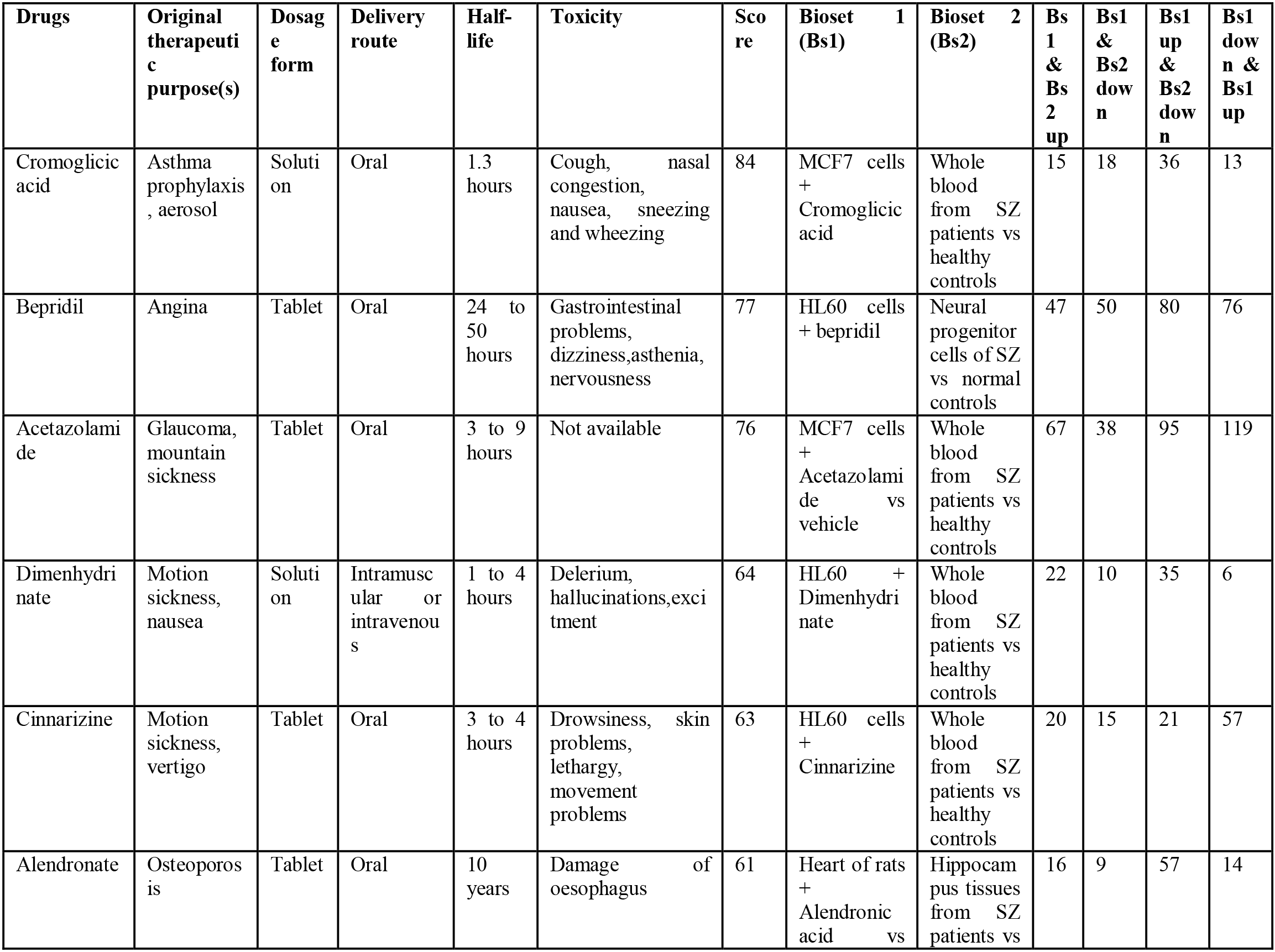
Details of the drugs that are identified as potentially repurposable for schizophrenia: Pharmacokinetic information is collected from DrugBank (www.drugbank.ca).

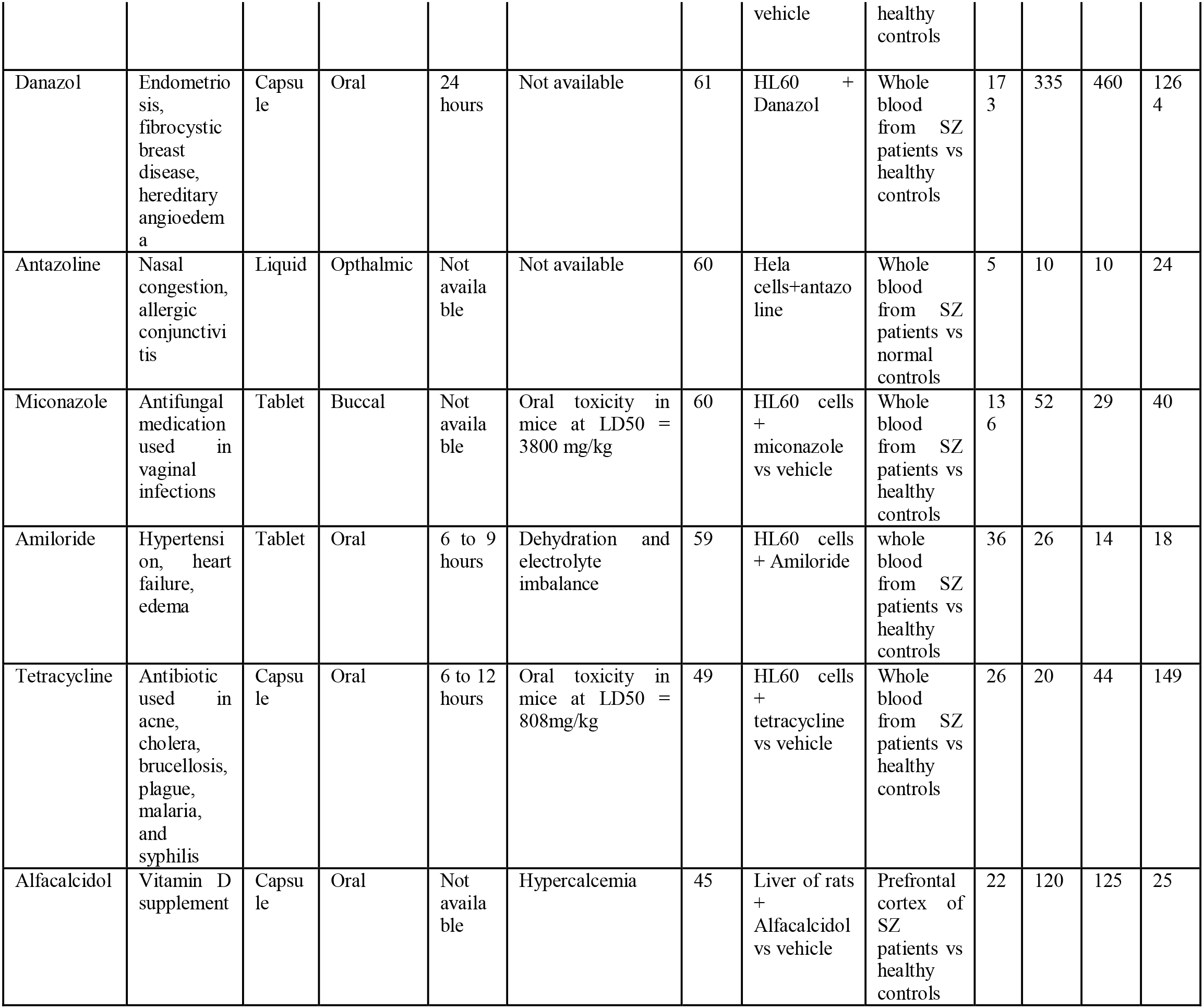

We queried the ClinicalTrials.gov database for these candidate drugs and found that cromoglicic acid and amiloride are being tested in clinical trials for their efficacy in Alzheimer’s disease and attention deficit hyperactivity disorder. It has been reported that cromoglicic acid in combination with ibuprofen reduces the levels of amyloid-beta protein levels, a pathological biomarker in Alzheimer’s disease, and promotes a neuroprotective state by activating microglia and inducing phagocytosis of amyloid-beta proteins.^15^ Minocycline, a broad spectrum tetracycline antibiotic, has been shown to be effective as an adjunctive drug, improving the effect of antipsychotic drugs in schizophrenia.^16^

### Literature based analysis

By surveying published literature, we found some evidence in support of these drugs for their use in schizophrenia. Alfacalcidol targets the protein VDR which was found to be overexpressed in whole blood obtained from schizophrenic patients compared to healthy controls (fold change (FC)=2.21, p-value=0.0037).^17^ Cinnarizine targets CACNA1H which is found to be overexpressed in neural progenitor cells differentiated for 2days from iPSCs (induced pluripotent stem cells) of schizophrenia patients versus healthy subjects (FC=3.1227, p-value=4.10E-20).^18^ We collected known and computationally predicted PPIs connecting the target genes of acetazolamide and queried the DisGeNet^19^ database whether any of the proteins in this network of interacting proteins are associated with neuropsychiatric disorders. Figure 1 shows only the protein targets from schizophrenia interactome. Here we considered all the proteins targeted by the shortlisted drugs. The protein targets of acetazolamide are carbonic anhydrases (CA*) and aquaporin (AQP1) (orange-colored nodes in Figure 2). The network of PPIs, including novel interactions, among the targets of acetazolamide shows that 19 genes are associated with various neuropsychiatric disorders: AQP1 and CA2, which are acetazolamide targets, DAXX, EPHB2, HSPD1, SLC4A3, SLC9A1, SRC, TCF4, TNK2, TRAF1, TRAF2, MTUS2, PICK1, GRM3, OLR1, TBP, PML and FOS (nodes with green border in Figure 2; Supplementary File 1), giving credence to the consideration that it has a potential application toschizophrenia. It has been shown to have high inhibitory activity against human CA II (hCA II), the ubiquitous cytosolic enzyme (inhibition constant, K_i_=12 nM), hCA VII (K_i_=2.5 nM), the brain-specific form of the enzyme, and mCA VII (K_i_=16nM).^20^ Human CA II (hCA II) and mouse CA VII (mCA VII) were found to be catalytically highly active (defined in terms of K_cat_/K_m_ for CO_2_ hydration described by two ionizations at pKa 6.2 and 7.5, with a maximum approaching 8×10^7^ M^−1^ s^−1^).^21^ K_cat_/K_m_ for hCA II is 1.5 ×10^8^ and for mCA VII, it is 7.6×10^7^.^21^ The increase in extracellular pH which accompanies neural activity is generated by the exchange of external H^+^ for cytosolic Ca^2+^. This process, and its impact on the glutamate receptors, NMDARs, has been shown to be regulated by CA14 in the synaptic microenvironment.^22^ On these lines, it is interesting to note that CA3 has been predicted to be a novel interactor of the glutamate receptor, GRM3, mutations in which have been associated with schizophrenia.^23^

**Figure 1.**
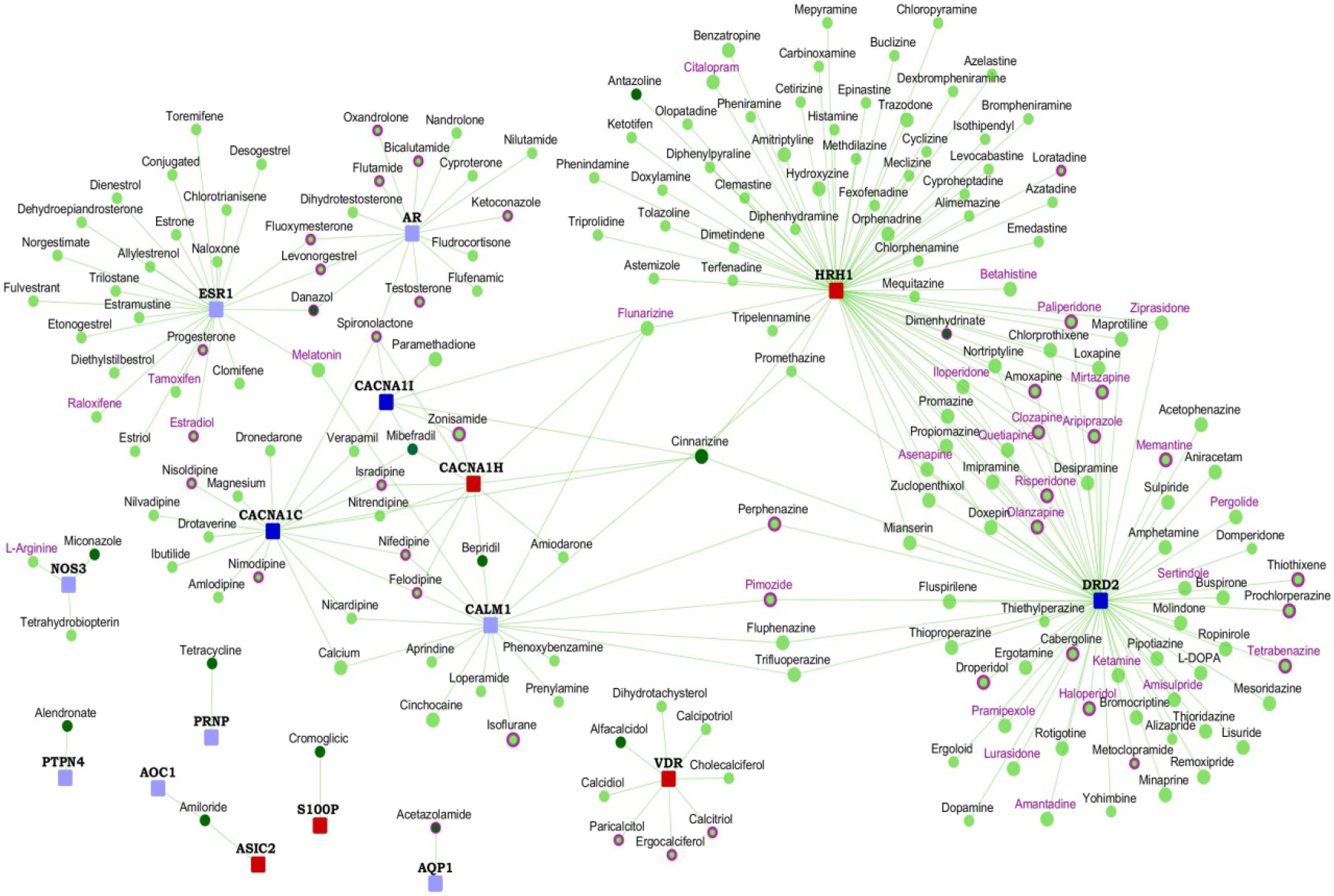
Drugs potentially repurposable for schizophrenia: The network shows the shortlisted drugs that may be potentially repurposed for schizophrenia. The shortlisted drugs are shown as round nodes colored in dark green, and other drugs are shown as light green nodes. FDA approved drugs are shown with purple borders. Drugs with purple labels are in clinical trials for schizophrenia. genes as square nodes colored in dark blue (schizophrenia genes), light blue (known interactors), and red (novel interactors). Schizophrenia genes are square nodes colored in dark blue, known interactors are colored in light blue, and novel interactors in red.

The protein targets of amiloride are ASIC1, ASIC2, AOC1, SLC9A1, PLAU, SCNN1A, SCNN1B, SCNN1G and SCNN1D (Figure 3). The network of PPIs among these targets of amiloride shows that 12 genes, including ASIC2, AOC1 and PLAU, which are amiloride targets, NEDD4, STX1A, MAPK1, HECW1, DAO, CSNK2A1, LASP1, SMG6 and PICK1are associated with various neuropsychiatric disorders. ASIC2 was a computationally predicted interactor of the gene SMG6, structural variants in which has been associated with schizophrenia or bipolar disorder in a Spanish population.^24,25^ SMG6 is located in the chromosomal region 17p13.3, linked to lissencephaly, a neuronal migration disorder arising from incomplete neuronal migration to the cerebral cortex during gestation, and characterized by an absence of normal convolutions in the cerebral cortex and an abnormally small head (or microcephaly).^25,26^ ASICs (acid-sensing channels) are members of the epithelial Na+ channel (ENaC) family of ion channels, expressed in the nervous system.^27^ It was shown in a study that ASIC2 is not expressed at the cell surface of high grade glioma (brain tumor) cells and this may be responsible for the constitutively activated inward Na^+^ current, which promotes increased cell growth and migration in these cells.^27^ In such glioma cells, compounds such as glycerol and the transcriptional regulator, sodium 4-phenylbutyrate, were shown to inhibit the constitutively activated inward Na^+^ current and reduce cell growth and migration.^27^ These compounds were shown to induce the movement of ASIC2 to the plasma membrane, and prevent the active inward current through negative regulatory mechanisms, reducing the ability of glioma cells to proliferate and migrate.^27^

**Figure 2.**
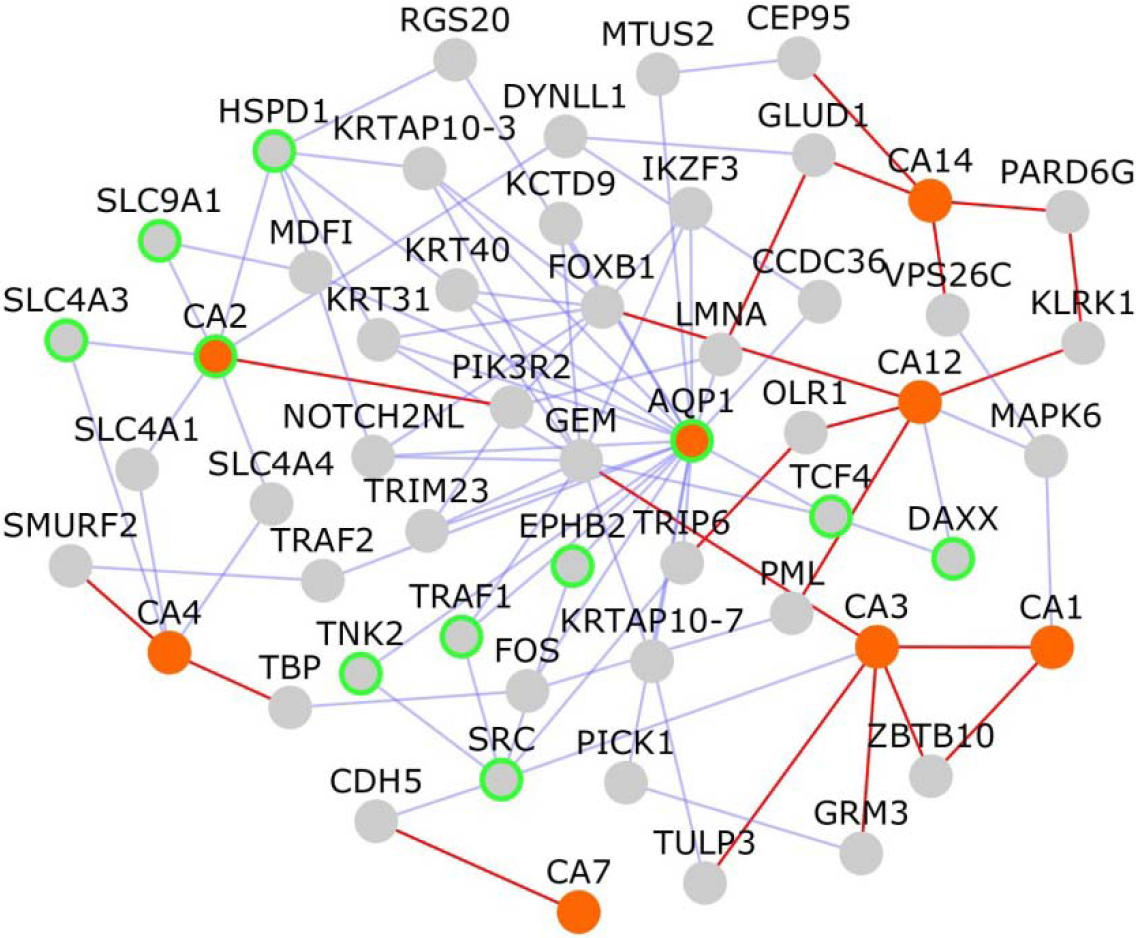
Network of PPIs among targets of acetazolamide: The network shows protein-protein interactions that connect the targets of acetazolamide, which are shown as orange colored nodes. Nodes that connect these target genes are shown as grey colored nodes. Nodes with light green borders are genes associated with neuropsychiatric disorders. Novel interactions are shown as red edges and known interactions as blue edges.

Both cinnarizine and antazoline target HRH1, which has been linked to schizophrenia etiology. Bacitracin has been shown to target A2M, linked to both Alzheimer’s disease and depressive disorder. Cinnarizine targets CACNA1C, associated with bipolar disorder, schizophrenia and depressive disorder, and CACNA1H, associated with epilepsy and autism. It targets DRD2, linked to bipolar disorder, schizophrenia, depressive disorder, Parkinson’s disease and attention deficit hyperactivity disorder. Danazol and miconazole target ESR1 and NOS3, both associated with Alzheimer’s disease. Tetracycline targets PRNP, linked to depressive disorder, Huntington disease-like 1 and Alzheimer’s disease.

We checked whether any of the genes having anti-correlated expression on cinnarizine treatment and in schizophrenia were associated with mammalian phenotype ontology (MPO) terms related to various morphological or physiological aspects of the nervous system http://www.informatics.jax.org/. It was found that mutations in 13 genes were associated with relevant MPO terms, namely,. AHI1, ENTPD1, IFNGR1, NAP1L1, NPTN, PIK3CA, PKN2, PRKDC, PTGS2, RBM12, SEC23A, SS18L1 and UBE3A. Two of these genes, IFNGR1 and AHI1, both linked to “abnormal depression-related behavior”, are predicted to have a novel interaction between them. Depressive symptoms have been observed in schizophrenia patients.^28^ IFNGR1 has been found to be necessary for the induction of IDO, the tryptophan synthesizing enzyme, which plays a role in depressive behavior, induced by inflammation.^28^ AHI1 is associated with susceptibility to schizophrenia and autism.^28^ Mice lacking neuronal expression of AHI1 had reduced levels of tyrosine kinase receptor B and a depressive phenotype, which was alleviated by antidepressants and overexpression of TRKB. ^28^ BDNF/TRKB signaling has been shown to play a key role in depression Altered BDNF/TRKB signaling in the prefrontal cortex, hippocampus and nucleus accumbens has been shown to give rise to depressive phenotype induced by inflammation.^29^ Another gene, UBE3A, was associated with increased dopamine level and serotonin level, abnormal brain wave pattern, cerebral cortex morphology, dendrite morphology, GABA-mediated receptor currents, long term potentiation and nervous system electrophysiology. Yet another gene, NPTN, was linked to abnormal synaptic transmission in the central nervous system and abnormal dendritic spine morphology.

**Figure 3.**
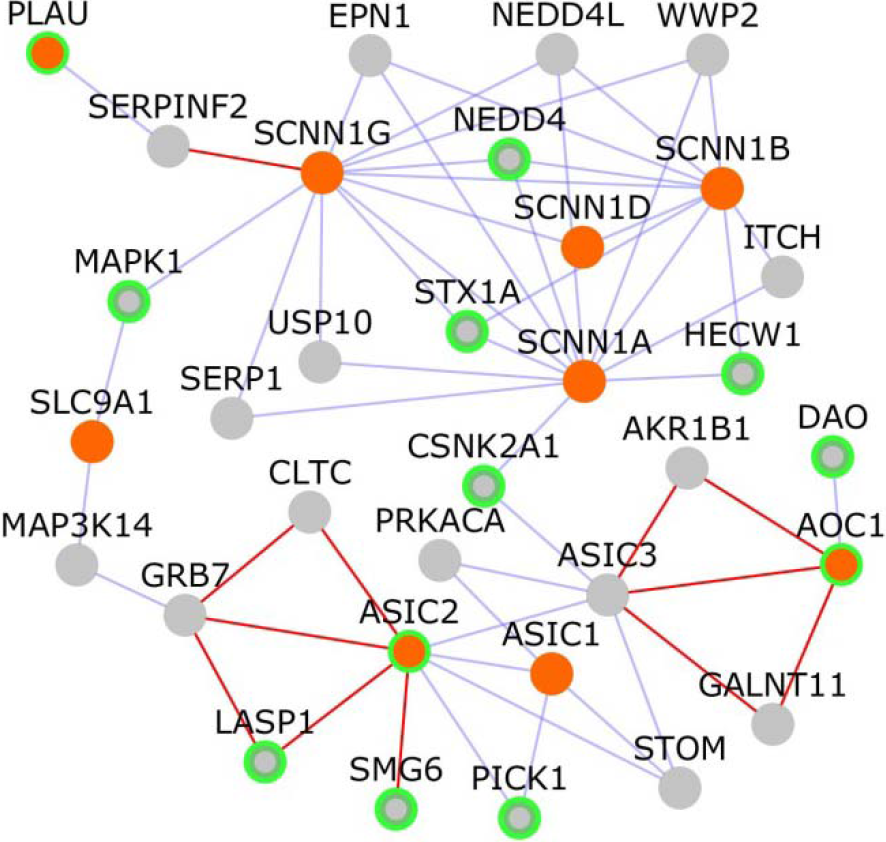
Network of PPIs among targets of amiloride: The network shows protein-protein interactions that connect the targets of amiloride, which are shown as orange colored nodes. Nodes that connect these target genes are shown as grey colored nodes. Nodes with light green borders are genes associated with neuropsychiatric disorders. Novel interactions are shown as red edges and known interactions as blue edges.

### Network analysis

We assembled the network of protein-protein interactions of genes that are differentially expressed by the shortlisted drugs and carried out network and enrichment analysis using a tool called LENS.^30^ The networks of genes found to be differentially expressed in acetazolamide, antazoline and cinnarizine, having an anti-correlation in schizophrenia, were shown to be enriched in ubiquitination and proteasome degradation pathways (Supplementary File 2). The ubiquitin proteasome system has been identified as an important pathway in several genetic studies of neuropsychiatric disorders including Alzheimer’s disease, Parkinson’s disease, psychosis and bipolar disorder.^31^ Many gene expression studies performed on blood collected from schizophrenia patients, and on post-mortem samples of hippocampus, prefrontal cortex and temporal cortex of patients have pointed at abnormalities in the ubiquitin proteasome pathway, which targets protein for degradation in the cell.^31^ Moreover, reduced protein ubiquitination, reduced levels of ubiquitin and ubiquitin-like activases and ligases, were identified in a region of the brain called the left superior temporal gyrus in schizophrenia patients.^31^ Left superior temporal gyrus, the volume of which has been shown to decrease in schizophrenia patients, is involved in the development of auditory hallucinations and thought process abnormalities seen in schizophrenia.^31^ Interestingly, acetazolamide which has been shown to mediate diuretic effects through its action on AQP1, induces AQP1 ubiquitination, and a proteasome inhibitor reversed its downregulatory action on AQP1.^32^ RAD51AP1 and AQR are novel interactors of the calcium channel CACNA1C and the nicotinic receptor CHRNA7 respectively in the schizophrenia interactome, found to have an anti-correlated expression in schizophrenia and acetazolamide treatment. It has been shown that UAF1, an interaction partner of USP1 deubiquitinating enzyme, associates with RAD51AP1, which interacts with RAD51 to mediate homologous recombination repair.^33^ NEDD4-1, an ubiquitin ligase, has been shown to promote the sorting of newly synthesized calcium voltage gated channels for proteasomal degradation.^34^ Suppression of AQR in HepG2, a liver cancer line, has been shown to inhibit protein ubiquitination.^35^ It has been shown that the expression of nicotinic receptors on the cell surface is regulated by the ubiquitin proteasomal system.^36^ The networks of genes found to be differentially expressed in alfacalcidol and tetracycline, having an anti-correlation in schizophrenia, were shown to be enriched in the neutrophil degranulation pathway (Supplementary File 2). Degranulating activity of neutrophils has been attributed to dysfunctional permeability of the blood-brain barrier in schizophrenia.^37^

Network of genes which were differentially expressed in acetazolamide and had an anti-correlation with schizophrenia were found to be significantly enriched in genes associated with rheumatoid arthritis (Supplementary File 2). Recently, the reduced prevalence of rheumatoid arthritis observed in schizophrenia patients was attributed to SNPs (single nucleotide polymorphisms) in the HLA region that conferred differential risk for schizophrenia and rheumatoid arthritis.^38^ The interactomes of schizophrenia and rheumatoid arthritis genes also showed a significant overlap and shared common pathways.^38^ The network of genes differentially expressed in alfacalcidol, cinnarizine and tetracycline were enriched in GWAS genes associated with inflammatory bowel disease (Supplementary File 2). The incidence of schizophrenia has been shown to be high in patients with immune-mediated inflammatory diseases such as inflammatory bowel disease, rheumatoid arthritis and multiple sclerosis.^39^ The network genes differentially expressed antazoline, with opposite expression in schizophrenia, were significantly enriched in GWAS genes associated with brain connectivity (Supplementary File 2). Abnormal interactions between brain networks have been pointed out to be an important contributing factor in schizophrenia etiology.^40^

We queried Drug Bank^41^ to find drugs that are structurally similar to the shortlisted drugs, or targeted the same genes as these drugs, and checked whether they demonstrated any clinical activity in schizophrenia or other neuropsychiatric disorders. Aviptadil, which is structurally similar to the shortlisted drug, bacitracin, showed neuroprotective effect in a mouse model of Parkinson’s disease by blocking microglial activation.^42^ Risperidone, nimodipine, nilvadipine, flunarizine, nifedipine, cannabidiol and clozapine target the same genes as cinnarizine. Flunarizine (targeting CALM1, CACNA1H) showed good efficacy and tolerability for the treatment of schizophrenia.^43^ Nifedipine (which targets CALM1, CACNA1H) enhanced learning and memory in schizophrenic patients with tardive dyskinesia.^44^ Cannabidiol (which targets CACNA1H) shows beneficial effects as an adjunctive drug along with existing anti-psychotic medication in schizophrenia.^45^ Risperidone (targeting DRD2) is used to treat schizophrenia, bipolar disorder, and irritability in autistic patients.^46–48^ Nimodipine (CACNA1C) has been found effective for treating resistant bipolar mood disorder.^49^ Nilvadipine (CACNA1C) was found to be effective in treatment of schizophrenia.^50^ Clozapine (targeting HRH1) is effective in treatment-resistant schizophrenia.^51^

## Discussion

We discuss here the potentially repurposable drugs, cinnarizine and alfacalcidol, for their biological characteristics.

Histamine receptors are highly expressed in brain regions associated with higher cognitive functions disturbed in schizophrenia.^52^ Leu49Ser mutation in HRH1 (*histamine receptor H1*) was associated with susceptibility to schizophrenia.^53^ Elevated levels of n-tele-methylhistamine, a histamine metabolite, was found in the cerebrospinal fluid of schizophrenia patents along with decreased HRH1 binding in frontal cortex and cingulate gyrus.^54^ Compounds acting at histamine receptors modulate extracellular striatal dopamine levels.^55^ The revised dopamine hypothesis of schizophrenia proposes that hyperactive dopamine transmission in the mesolimbic areas such as the ventral tegmental area and ventral striatum including nucleus accumbens may contribute to disease etiology.^56^ Enhanced release of neuronal histamine was observed on DRD2 (*dopamine receptor D2*) activation and in methamphetamine/phencyclidine-induced animal models of schizophrenia.^57,58^ Histamine antagonists inhibit behavioral sensitization arising from increased levels of extracellular dopamine.^58–61^ The fact that refractory schizophrenia may be treated with clozapine, an HRH1 antagonist, indicates that extra-dopaminergic systems, viz. the histamine neuron system, contribute to schizophrenia etiology.^51,58^ Clozapine also has strong affinity to dopaminergic receptors and decreases hyperactivity of the mesolimbic dopaminergic pathway by blocking 5-HT2A (*5-hydroxytryptamine receptor 2A*).^56^ Famotidine, an HRH2 antagonist, significantly reduced psychotic symptoms in schizophrenia patients.^62^ The examples of clopazine and famotidine indicate that a drug acting as a DRD2 and HRH1 antagonist may serve to alleviate psychotic symptoms arising from the interplay of dopaminergic and histamine neuron systems. Cinnarizine, an HRH1, DRD2 and Calcium channel antagonist commonly used to treat motion sickness, may be re-purposed to treat symptoms of schizophrenia (see Figure 4).^63^ It prevents vesicular uptake of dopamine.^64^ It shows antagonistic activity at the Calcium channel, CACNA1C, whose reduced levels attenuate the function of the mesolimbic dopaminergic pathway and impair behavioral responses to dopamine stimulants.^65^ Calcium channel antagonists reduce neurotransmission of dopamine.^66^ HRH1 was predicted to interact with the schizophrenia gene NAB2. NAB2 modifies the induction of DARPP-32, which modulates the response to dopamine in striatal neurons.^67^ HRH1 has also been predicted to interact with BBS5, a ciliary protein. BBS5 interacts with DRD1 and is involved in translocating DRD1 out of the cilia in response to dopamine receptor agonists, thereby implicating neuronal cilia in dopamine signaling.^68^ BBS5 was predicted to interact with SLC6A15, which is enriched in striatal DRD2 neurons and inhibited by loratadine, an HRH1 antagonist.^69,70^

**Figure 5.**
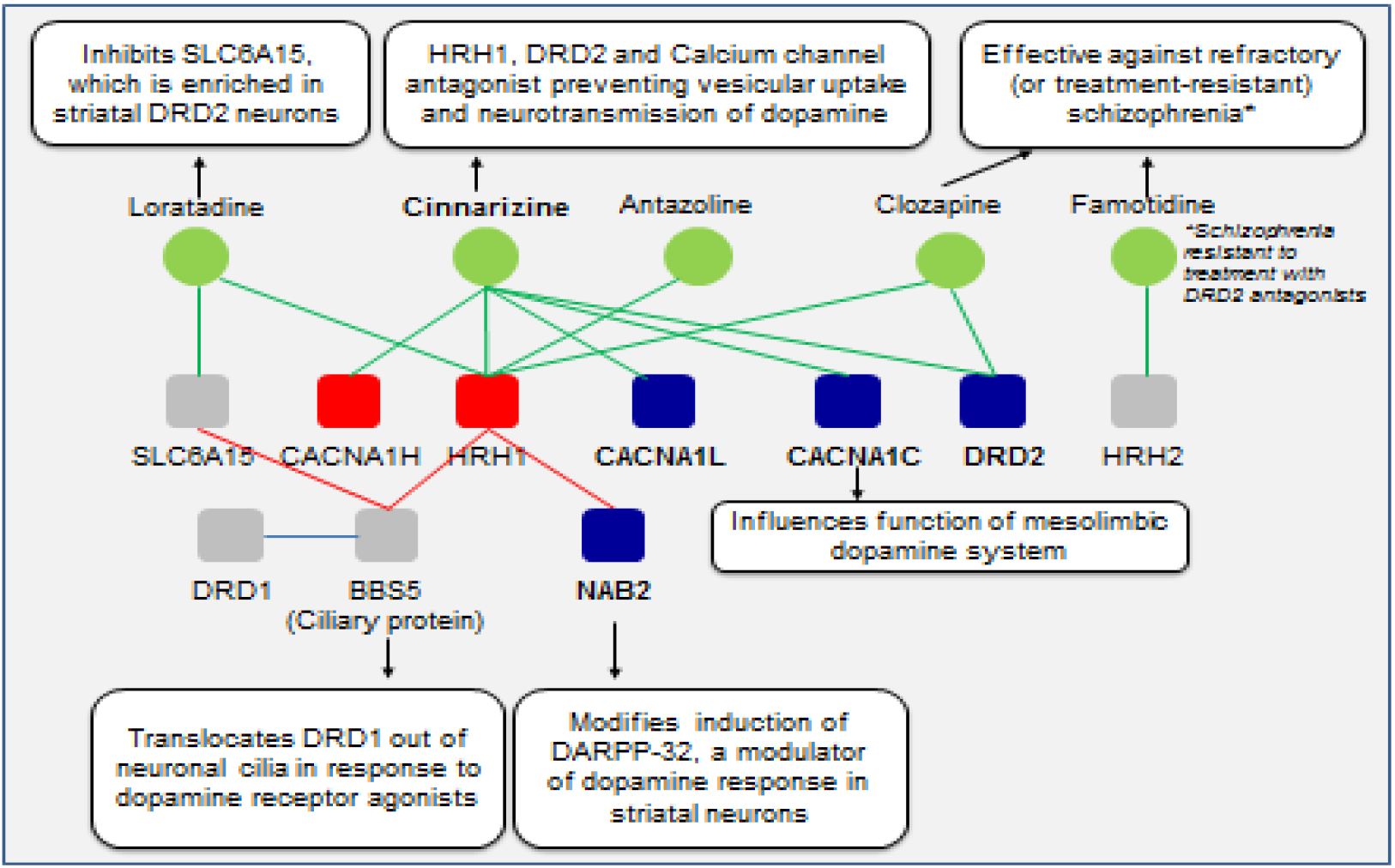
Cinnarizine and its targets in the schizophrenia interactome: The drug cinnarizine is shown here with the proteins it targets from the schizophrenia interactome. 4 additional proteins (BBS5, DRD1, HRH2 and SLC6A15) and 3 additional drugs (loratadine, clozapine and famotidine) that are relevant to the hypothesis are also shown. Cinnarizine, targets 3 schizophrenia genes and two novel interactors which constitute calcium channels, and histamine & dopamine receptors. Since histamine antagonists are known to reduce dopamine levels through their action on dopamine receptors, and calcium channel antagonists are known to reduce dopamine neurotransmission, the HRH1, DRD2 and calcium channel antagonist, cinnarizine, may be repurposable for schizophrenia. Another shortlisted drug, antazoline, is not part of the reasoning presented here even though it is an HRH1 antagonist. Schizophrenia genes are shown as dark blue colored nodes, novel interactors are red colored nodes and genes relevant to the hypothesis, which are not in the

Deficiency of vitamin D which exerts its effects through VDR (*vitamin D receptor*) has been observed in schizophrenia patients.^71^ Interplay between the vitamin D system and the dopaminergic system, which plays a key role in schizophrenia etiology, has also been noted.^72^ VDR is highly expressed in regions of the brain associated with schizophrenia, viz. hippocampus, prefrontal cortex and dopaminergic neurons in substantia nigra of rats and humans.^73^ During early stages of development, VDR is expressed in the mesencephalon precisely at the time when monoamine cells differentiate to dopaminergic cells and dopaminergic systems are innervated.^72^ Reduced levels of the enzyme COMT (*catechol-O-methyltransferasS*), which converts the dopamine metabolite DOPAC (*3,4-Dihydroxyphenylacetic acid*) into HVA (*homovanillic acid*) and affects dopamine turnover, have been observed in mice pups having Vitamin D deficiency.^72^ In rats with vitamin D deficiency, the effect of MK-801, an NMDA (*N-methyl-D-aspartate*) receptor antagonist which indirectly activates dopaminergic activity and also induces hyperlocomotion in animals, was found to be attenuated with the use of haloperidol, a DRD2 (*Dopamine Receptor D2*) anatgonist.^74^ In SH-SY5Y cells routinely used to model neural functions, overexpression of VDR resulted in increased levels of dopamine, overexpression of TH (*tyrosine hydroxylase*) which is an enzyme involved in the production of the precursor of dopamine called L-DOPA and overexpression of DRD2 whose increased activity has been noted in models of schizophrenia, among other regulatory effects on genes associated with the dopaminergic system.^75,76^ On treatment of these SH-SY5Y cells with calcitriol, a biologically active form of vitamin D, increased levels of dopamine metabolites such as HVA, increased COMT levels and reduced DRD2 expression were observed.^76,77^ Hence, dopaminergic aspects of schizophrenia etiology as proposed by the dopamine hypothesis of schizophrenia may be, at least in part, treated by vitamin D supplementation. ^56^ In this respect, vitamin D supplementation in the first year of life has been associated with reduced risk of schizophrenia in males, in a study based on 9114 subjects from Northern Finland 1966 birth cohort.^78^ The drug alfacalcidol, an analog of vitamin D, commonly used as a vitamin D supplement, or to treat conditions involving imbalance in calcium metabolism such as hypercalcemia and imbalance in bone metabolism such as osteoporosis, may be potentially re-purposed to treat dopaminergic symptoms in schizophrenia, possibly in combination with dopamine receptor antagonists such as clozapine.^79,80^ Alfacalcidol acts on VDR predicted to interact with the schizophrenia gene STAG1 in the schizophrenia interactome. STAG1 (*Stromal Antigen 1*) is a subunit of the cohesion complex required for cohesion of sister chromatids after DNA replication, possibly also playing a role in role in spindle assembly during cell division.^81^ Even though STAG1 has been significantly associated with schizophrenia, its contribution to disease etiology remains elusive.^82^ However, STAG1 has been found to be differentially expressed in a neuronal progenitor cell line derived from the developing cerebral cortex in which TCF4, a transcriptional regulator having common variants significantly associated with schizophrenia, was underexpressed.^83–85^

TCF4 (*transcription factor 4*) is a schizophrenia gene. Apart from its involvement in brain development, synaptogenesis, cell cycle exit, cell proliferation and differentiation, TCF4 has been reported to be an indirect target of the VDR pathway in mammary gland of mice and in cell lines of colorectal cancer.^86^ TCF4 has also been described as a negative modulator of oligodendrocyte maturation, necessary for the process of formation of the fatty cover that insulates axons in neurons called myelination in the brain.^87^ It has been found that while myelin loss accompanied by a decrease in the number of cells from the oligodendrocyte lineage expressing TCF4 induced by chronic stress, can be reversed by treatment with lithium (used in the treatment of major depressive disorder), mice lacking DRD2 which display depression-like behavior and has reduced myelin levels compared to wild type are not sensitive to lithium treatment.^88,89^ This shows that some pathway involving TCF4 which is necessary for re-myelination has been disrupted in mice lacking DRD2.^88^ Within purview of the fact that defects in myelination is a pathological marker of schizophrenia, this is interesting because two rare mutations (G428V and P299S) found in TCF4 that were associated with schizophrenia have been found to have increased transcriptional activity of TCF4 and in another study, density of mature oligodendrocytes essential for myelination have been observed to be reduced in white matter derived from prefrontal cortex of schizophrenia patients, which is an expected outcome if TCF4, a negative modulator of mature oligodendrocytes were to be overexpressed.^90–92^ Also, treatment with DRD2/DRD3 agonist, quinpirole, has shown a decrease in the number of mature oligodendrocytes while treatment with haloperidol, a DRD2 antagonist, has shown an increase in the number of mature oligodendrocytes.^93^ Many studies have noted this interplay of myelination and dopaminergic systems.^94^ GDNF (*glial cell derived neurotrophic factor*), which is expressed in oligodendrocytes, plays a critical role in the survival of dopaminergic neurons and polymorphisms in GDNF have been associated with schizophrenia.^95,96^ Moreover, it has been reported that an upregulation in DRD2 signaling led to increased expression of GDNF.^97^ Increased dopamine levels in the prefrontal cortex and impairment in behavioral responses related to dopamine were observed in mice treated with cuprizone, a drug that induces de-myelination. ^94^ It has been observed that psychiatric symptoms which may be treated with clozapine, a DRD2 antagonist, co-occur in multiple sclerosis. ^94^ Multiple sclerosis is a disease in which myelin sheath which insulates the brain and the spinal cord is impaired resulting in de-myelination.^98^ It is interesting to note that oligodendrocyte maturation and myelination are disrupted in multiple sclerosis lesions, when VDRs which are abundantly present on oligodendrocyte progenitor cells are blocked.^99^ On the other hand, oligodendrocyte maturation and re-myelination are observed when VDRs are activated by vitamin D.^99^ In a clinical study, the drug alfacalcidol was shown to be effective for treatment of fatigue experienced by multiple sclerosis patients and it has been proposed that fatigue in multiple sclerosis may be related to impaired dopaminergic system.^100,101^ Increased fatigue in multiple sclerosis has been associated with reduced integrity of white matter in ventromedial prefrontal cortex which is innervated by dopaminergic projections.^101^ Impaired myelin formation manifesting as reduced density of processes on oligodendrocytes was found in cortical tissues of stillborn affected with Cornelia de Lange syndrome, a syndrome characterized by distinctive facial features, behavioral abnormalities and limb deformations, and associated with defects in the cohesion complex constituting STAG1. ^102,103^ Hence, the predicted interaction of VDR with STAG1 and its association with TCF4 may show that re-purposing alfacalcidol for schizophrenia in combination with dopamine receptor antagonists may potentially alleviate dopaminergic symptoms and myelin defects influenced by the dopaminergic system in schizophrenia, when administered, perhaps, at early stages of the disease.

## Methods

### Identification of potentially repurposable drugs using NextBio

We first constructed protein-protein interaction network of schizophrenia genes, and then identified the drugs that target any of the proteins in this interactome. Several of these drugs were known to have therapeutic value for nervous system, but there were several drugs that targeted other anatomical systems in the human body. As a mechanism of shortlisting drugs for further analysis, we selected those that targeted multiple proteins of the schizophrenia interactome, novel protein interactors, or those that target proteins that are also targeted by many drugs. Next step involved identifying the drugs that have opposite differential expression to the differential expression of schizophrenia (i.e., genes over-expressed in schizophrenia are under-expressed by drug treatment and vice versa). We studied each of these drugs in comparison to gene expression profiles of schizophrenia by using the software suite NextBio (now called BioSpace). NextBio is used to study the effect of diseases and/or drugs on publicly available gene expression data.^104^ Bioset 1 (‘BS1’) or a particular cell line in which differential expression by drug has been studied was compared with a bioset 2 (‘BS2’), a cell lines in which differential expression in schizophrenia patients was studied. A correlaton score is generated by the tool based on the strength of the overlap or enrichment, between the two biosets. Additional statistical criteria such as correction for multiple hypothesis testing are applied and the correlated biosets are then ranked by statistical significance. A numerical score of 100 is assigned to the most significant result, and the scores of the other results are normalized with respect to the top-ranked result. We excluded drugs with unacceptable levels of toxicity or undesirable pharmacokinetics.

### Network analysis using LENS

LENS (Lens for Enrichment and Network Studies of human proteins) is a web-based tool which may be used to identify pathways and diseases that are significantly enriched among the genes submitted by users.^30^ The LENS algorithm finds the nearest neighbor of each gene in the interactome and includes the intermediate interactions that connect them. LENS then computes the statistical significance of the overlap of genes in the network and genes with annotations pertaining to pathways, diseases, drugs and GWASs, and reports a P-value computed from Fisher’s exact test.

Shortlisted drugs which are being tested in clinical trials against various neuropsychiatric disorders were identified from NIH Clinical Trials (https://clinicaltrials.gov/).

Differential expression of the novel interactor VDR in whole blood obtained from schizophrenia patients was identified from GSE38485^17^, and that of CACNA1H in iPSCs of schizophrenia patients was identified from GSE92874^18^.

Association of the various genes in the network of PPIs among targets of the shortlisted drugs was identified from DisGeNET, a database that integrates human gene-disease associations from expert curated databases and text-mining derived associations.^19^

Drugs that are structurally similar to the shortlisted drugs, or targeted the same genes as these drugs were identified from Drug Bank (https://www.drugbank.ca/).^41^

## Author contributions

The study has been designed by MKG and in part by KBK. SC carried out correlation analysis of drugs against diseases. KBK carried out literature study and further bioinformatics analysis of selected drugs. Manuscript has been prepared by KBK and edited by MKG.

## Acknowledgements

This work has been funded by the Biobehavioral Research Awards for Innovative New Scientists (BRAINS) grant R01MH094564 awarded to MKG by the National Institute of Mental Health of National Institutes of Health (NIMH/NIH) of USA. The content is solely the responsibility of the authors and does not necessarily represent the official views of the National Institute of Mental Health, the National Institutes of Health. MKG thanks Ansuman Chattopadhyay of University of Pittsburgh Health Sciences Library System for licensing and demonstration of molecular biology software resources.

## Supplementary files

**Supplementary file 1 – Genes associated with neuropsychiatric disorders**: Neuropsychiatric disorders associated with genes in the network of PPIs of acetazolamide and amiloride targets are shown in this file

**Supplementary file 2 – GWAS traits, diseases and pathways from LENS analysis**: GWAS traits, diseases and pathways enriched in the network of PPIs of genes found to have anti-correlated expression in schizophrenia and on administration of the potentially repurposable drugs are shown in this file

